# Vessel wall reinforcement metrics as drought resistance indicators in angiosperm fossil wood assemblages

**DOI:** 10.1101/2022.11.08.515713

**Authors:** Hugo I. Martínez-Cabrera, Emilio Estrada-Ruiz, Carlos Castañeda-Posadas

## Abstract

**Background:** Plant ecologists have developed methods to measure xylem drought resistance but these cannot be used in fossil woods. There is, however, one anatomical trait highly correlated with cavitation resistance: the squared vessel-wall thickness-to-span ratio ((t/b)^2^_h_). This metric though, could be in many cases impractical to measure in fossil samples because they often are small and sample sizes are seldom reached.

**Questions:** are there alternative anatomical metrics that could be used instead of (t/b)^2^_h_ to infer drought resistance of fossil wood assemblages?

**Study site and dates:** 279 species belonging to 14 extant communities from North and South America. Three fossil wood floras from the Oligocene and Miocene of Mexico.

**Methods:** We calculated three alternative wall reinforcement metrics to determine their relationship with (t/b)^2^_h_ and drought resistance. These are based on vessel diameter and vessel wall thickness.

**Results:** We found that one of the alternative metrics ((t/b)^2^_hydraulic mean_) could potentially be used instead of (t/b)^2^_h_. The widely measured vessel wall-to-lumen ratio (VWLR), was the closest related to climate, and thus helpful in identifying broad precipitation differences among floras. VWLR and (t/b)^2^_h_ might be describing slightly different ecological axes of ecological variation, with the latter associated with investment in support tissue, in addition to water availability alone.

**Conclusions:** Some of the alternative metrics we explored can be used, in combination with other functional traits, to better describe fossil forest functional strategies.

## Introduction

Because the diffusion coefficient of water is larger than for CO_2_ (Lambers *et al*., 1998), carbon fixation could be an expensive process in terms of water expenditure. Per each mole of CO_2_ fixed during photosynthesis, over 100 moles of water are transpired (Cramer *et al*. 2008). Efficient carbon fixation therefore requires an equally efficient hydraulic system to meet the water requirements of the evaporative surface during the photosynthesis. For this reason, stem hydraulic capacity is directly related to growth rate (e.g. Machado & Tyree 1994) and photosynthetic capacity (Santiago *et al*. 2004). There is a general relationship between vessel diameter (the main determinant of xylem hydraulic efficiency, Poorter 2008, Zanne *et al*. 2010) and environmental conditions such that, across vegetation types (in low latitudes and altitudes), water availability and vessel size are positively related (Carlquist 1988, Wheeler *et al*. 2007). In wet warm environments, plants maximize hydraulic efficiency by decreasing water flow resistance in the xylem (i.e. by having larger vessel diameter) to maintain high transpiration rates, carbon fixation and growth (Tyree 2003). On the other hand, in dry environments these large, efficient vessels are at disadvantage since they are more likely to experience drought induced cavitation (because the likelihood of finding large pores in the pit membrane increases with vessel size; Wheeler *et al*. 2005) and/or their walls are more likely to experience mechanical failure (Hacke & Sperry 2001, Hacke *et al*. 2001, Jacobsen *et al*. 2005).

Plants from drier regions are generally more able to cope with drought because cavitation in small vessels occurs at lower water potential than in wet adapted plants with larger vessel diameters (Sperry & Pockman 1993, Kolb & Sperry 1999). The ecological outcome of vessel cavitation is the disruption of the water column, which results in the reduction of plant water supply to leaves (Meinzer *et al*. 2001), and a consequential decrease in stomatal conductance (Pratt *et al*. 2005) and photosynthesis (Brodribb & Feild 2000). Knowing how these wood anatomical traits related to drought resistance vary in modern ecosystems, can help us to infer some aspects of fossil forest function, particularly of those traits associated with adaptations to water deficit.

In living plants, cavitation resistance is measured using the water potential at which there is a determined percent loss of hydraulic conductivity. This percent is usually 50% (P_50_). As P_50_ cannot be directly measured in fossil woods, the squared vessel-wall thickness-to-span ratio ((t/b)^2^_h_) which is tightly related with P_50_ (Hacke *et al*. 2001), has been used to infer its values in a fossil wood assemblage (Martínez-Cabrera & Estrada-Ruiz 2014). (t/b)^2^_h_ is a good indicator of resistance to water stress because it explains up to 95% of the variation in P_50_ (Jacobsen *et al*. 2005). (t/b)^2^_h_ is, however, hard to measure because it only considers those vessel pairs that fall within 3 to 5 μm of the hydraulic diameter. Because the small size of many fossil woods, it is common to find very small sample sizes with no more than a couple of vessel pairs falling within 5 μm of the mean hydraulic diameter. Besides, many fossil assemblages have species with exclusively solitary vessels, rendering this metric impractical. Given the promising results of the vessel wall reinforcement in providing ecological information in wood paleofloras (Martínez-Cabrera & Estrada-Ruiz 2014), here we explored if more easy-to-measure metrics are equally informative. Specifically, 1) we used anatomical data from extant communities to quantify how much of the variation of the squared vessel-wall thickness-to-span ratio is explained by three alternative metrics, and if one of these could be used to infer drought resistance. In addition, 2) we correlated these alternative vessel wall reinforcement metrics with climate variables to if they are likely to provide information about the growing conditions and functional characteristics of fossil floras. Finally, 3) we calculated the metric with tighter relation with climate variables (vessel wall-to-lumen ratio) of three relatively well known fossil wood assemblages, to contrast with the functional and climate interpretations provided by other studies in those sites.

## Material and methods

We analyzed two different databases (Table S1). With the fist database, that includes 62 species from a variety of environments in North and South America (Martínez-Cabrera *et al*. 2009), we tested if (t/b)^2^_h_ variation is mirrored by more simple versions of it, such as (t/b)^2^_mean_, (t/b)^2^_hydraulic mean_, and VWLRT. The squared vessel-wall thickness-to-span ratio (t/b)^2^_h_ (where *t*, is the thickness of the double-wall between two adjacent vessels and *b* is the diameter of the conduit closest to the hydraulic mean diameter), is exclusively measured in vessel pairs within 3 to 5 μm of the hydraulic diameter. The mean hydraulic diameter has to be determined before locating the target vessel pairs. In the alternative metrics we explored here ((t/b)^2^_mean_ and (t/b)^2^_hydraulic mean_), *t* is the mean of vessel wall thickness times two, while *b* is simply the mean vessel diameter and the mean hydraulic diameter of the sample, respectively. The difference between (t/b)^2^_h_ and (t/b)^2^_hydraulic mean_ is that in the former, *t* only includes the vessel pairs falling within 5 μm of the mean hydraulic diameter, while in the latter, *t* is simply the mean hydraulic diameter of the sample. In this database, vessel lumen diameter, including mean vessel diameter and hydraulic mean was calculated using diameters of circles with the same area as the individual vessel lumens. The hydraulic mean was calculated as the sum of the contribution of all conduit diameters (∑*d^5^*) divided by the total number of vessels (∑*d^4^*) (see Martínez-Cabrera & Estrada-Ruiz 2014 for details). VWLR is the ratio between wall thickness and the mean vessel diameter. We calculated the P_50_ values (based on the general formula P_50_=20.662–154.646x, where x was (t/b)^2^_h_, (t/b)^2^_mean_ or (t/b)^2^_hydraulic mean_) for each one of the three metrics and compared the estimates to assess the suitability of (t/b)^2^_mean_ and (t/b)^2^_hydraulic mean_, as an alternative to the P_50_ values predicted by (t/b)^2^_h_. The original formula was derived using (t/b)^2^_h_ (formula provided by Uwe Hacke).

For our second analysis, we then combined the climate (MAP, MAT, and PET) and VWLR data from the database mentioned above (Martínez Cabrera *et al*. 2009) with the second database (Martínez-Cabrera & Cevallos-Ferriz 2008). This database comprises extant floras growing under a wide range of environments and was used to determine the extent to which VWLR is environmentally driven. As (t/b)^2^_h_ was not measured in that study (Martínez-Cabrera & Cevallos-Ferriz 2008), it was not analyzed here. Lastly, we analyzed the extant communities’ VWLR with that of three fossil localities, El Cien Formation (Oligocene-Miocene; Martínez-Cabrera & Cevallos-Ferriz 2008), Las Guacamayas (Miocene, Chiapas, Mexico) and La Mina (Miocene, Tlaxcala, Mexico) (Castañeda-Posadas 2007) to establish the scope and resolution of the paleoecological information that this anatomical trait provides. We did not have the mean hydraulic diameter for the fossil woods, and therefore it was not possible to estimate P_50_ values based on (t/b)^2^_hydraulic mean_, and decided not to use the mean vessel diameter because, as we discuss below, the P_50_ calculated with (t/b)^2^_mean_ has larger associated errors.

## Results and Discussion

### Relationship between the squared vessel-wall thickness-to-span ratio and alternative metrics

As expected, (t/b)^2^_h_ was more tightly related with (t/b)^2^_hydraulic mean_ (R^2^ = 0.83, P < 0.001), than to (t/b)^2^_mean_ (R^2^ = 0.54, P < 0.001) or VWLR (R^2^ = 0.54, P <0.001) (Figure 1). Consequently, the P_50_ values based on (t/b)^2^_h_ are more similar to those calculated using (t/b)^2^_hydraulic mean_, (R^2^ = 0.83, P < 0.001; Figure 2A) than those based on (t/b)^2^_mean_ (R^2^ = 0.54, P < 0.001). (t/b)^2^_mean_ tends to overestimate P_50_ values (cavitation at more negative water potentials), especially in drier communities (Figure 2B), but this is not the case for (t/b)^2^_hydraulic mean_. Our results suggest that if (t/b)^2^_h_ is not possible to measure in a particular fossil wood assemblage, (t/b)^2^_hydraulic mean_ could be used to broadly infer cavitation resistance. We suggest, however, that comparing P_50_ estimates of fossil wood assemblages calculated with different metrics (*i.e.* P_50_ vs. P_50_ hydraulic) should be avoided as the estimated error varies (Figure 2).

**Figure 1.**
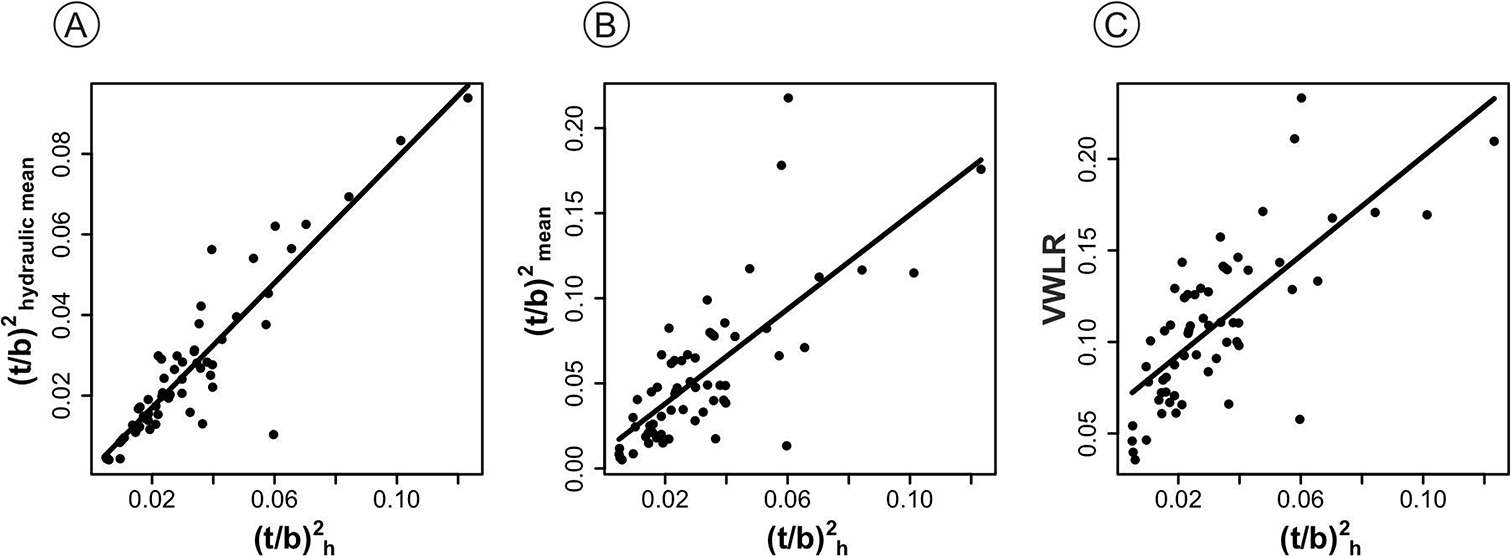
Relationship between the squared vessel-wall thickness-to-span ratio (t/b)^2^_h_ and A. (t/b)^2^_hydraulic mean_. B. (t/b)^2^_mean_. C. VWLR.

**Figure 2.**
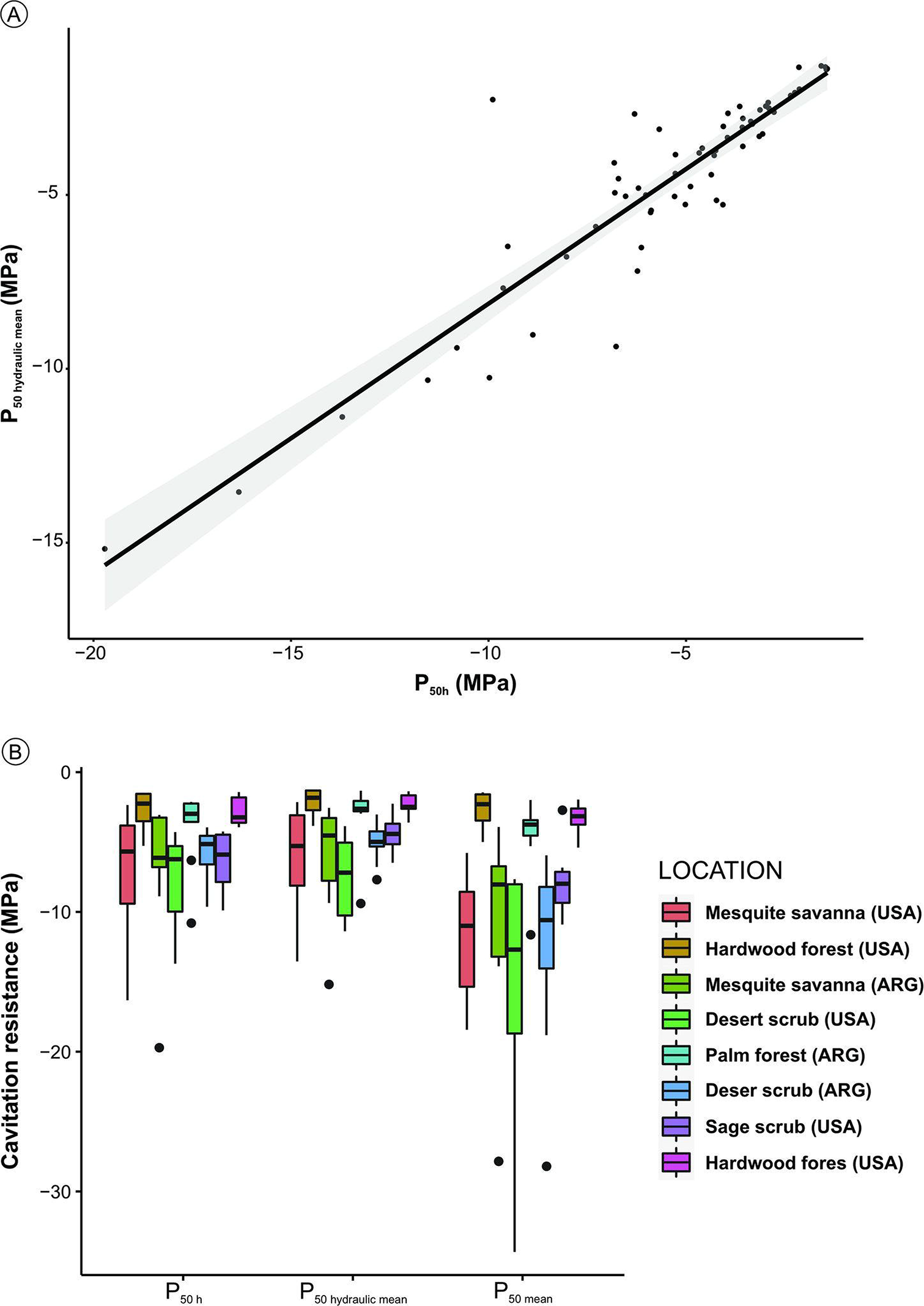
A. Relationship between (t/b)^2^_h_ and (t/b)^2^_hydraulic mean_ predicted P_50_ values. B. Boxplot showing predicted P_50_ values using the original the squared vessel-wall thickness-to-span ratio metric (P_50h_), the mean hydraulic diameter (P_50hydraulic mean_), and mean vessel diameter (P_50mean_). Boxplots show median, interquartile range and largest values within 1.5.times the interquartile range below the 25^th^ and above the 75^th^ percentiles.

### Relationship between wall reinforcement metrics climate

In general, vessel reinforcement (all four metrics) increases with temperature (MAT) and potential evapotranspiration (PET) and decreases with precipitation (MAP). MAP was the climate variable that better explained variation of all four-vessel reinforcement metrics (Figure 3A), but its relationship with VWLR was the tightest. VWLR was the anatomical trait more closely related to all climate variables (Figure 3, Table 1). In this sense, (t/b)^2^_h_ and VWLR seem to be describing a slightly different ecological axis of variation. Based on our results, VWLR has more potential as a tool in paleoclimate prediction, while (t/b)^2^_h_ is describing, in addition to climate, cavitation resistance and investment in support, and can thus can offer a supplementary layer of paleoecological information. Although fiber characteristics are the main determinants of investment in support (*i.e.,* wood density, Hacke *et al*. 2001), the link between wood density and (t/b)^2^_h_ is via a coordinated variation between fiber and vessel wall-to-lumen ratios (Hacke *et al*. 2001). Jacobsen *et al*. (2005) proposed that the link between wood density and (t/b)^2^_h_ is through the indirect effect of fibers reinforcing vessel walls, since higher (t/b)^2^_h,_ in their study, was not associated with a decrease conduction efficiency.

**Figure 3.**
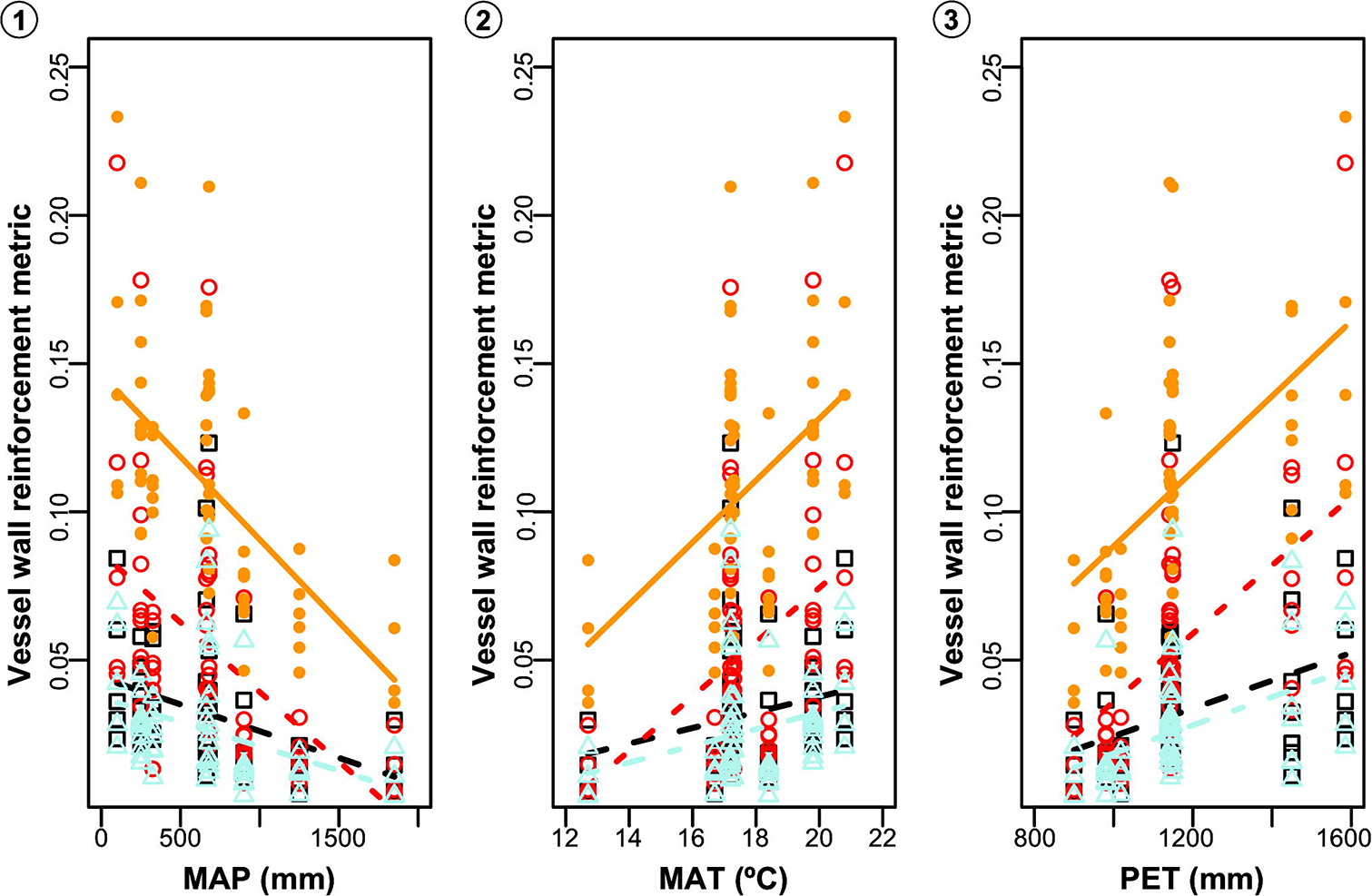
Climate variables as a function of vessel wall reinforcement metrics. A. MAP. B. MAT. C. PET. The squared vessel-wall thickness-to-span ratio ((t/b)^2^_h_) black squares and dashed line; (t/b)^2^_mean_ red rings and dotted line; (t/b)^2^_hydraulic mean_ light blue triangles and dot-dashed line; VWLR solid orange circles and continuous line.

**Table 1.**
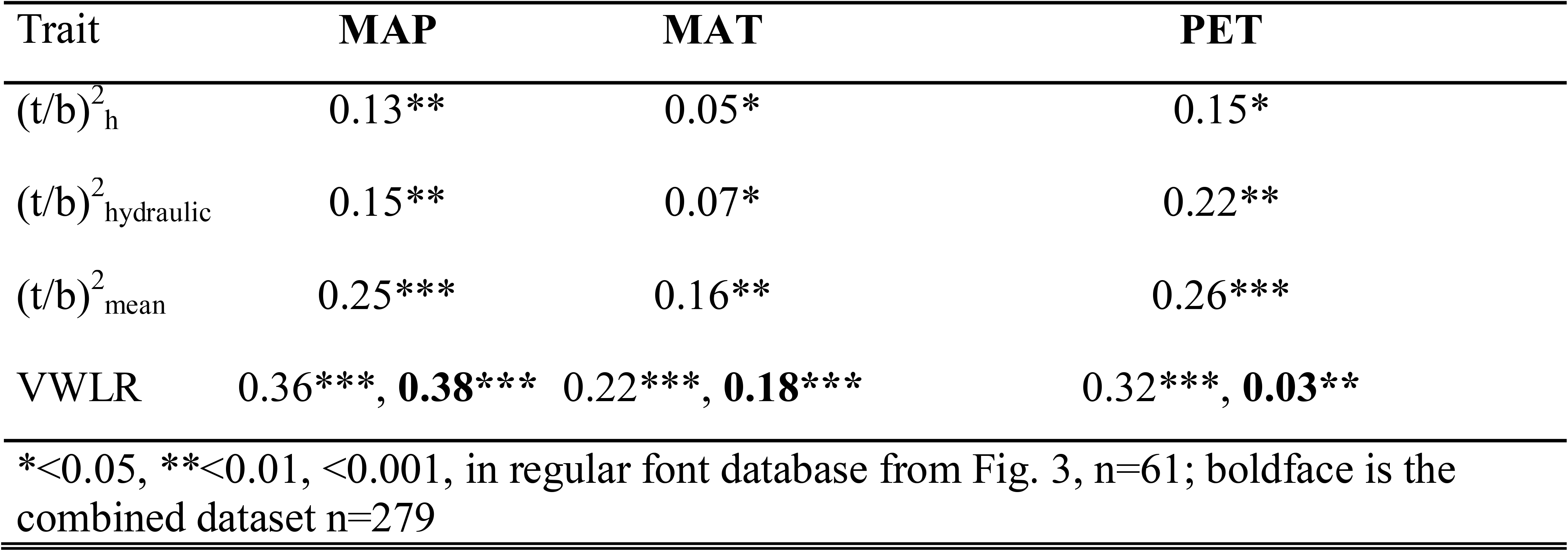
Regression coefficients for the relationship between climate variables and vessel wall reinforcement metric.

| Trait | MAP | MAT | PET |
| --- | --- | --- | --- |
| $(t/b)^2_h$ | 0.13** | 0.05* | 0.15* |
| $(t/b)^2_{hydraulic}$ | 0.15** | 0.07* | 0.22** |
| $(t/b)^2_{mean}$ | 0.25*** | 0.16** | 0.26*** |
| VWLR | 0.36***, **0.38\*\*\*** | 0.22***, **0.18\*\*\*** | 0.32***, **0.03\*\*** |
| *$<0.05$, **$<0.01$, ***$<0.001$, in regular font database from Fig. 3, n=61; boldface is the combined dataset n=279 | | | |

However, (t/b)^2^_h_ is not always correlated with the proportion of fibers surrounding vessels (*i.e.,* species with extremely dense wood might have vasicentric parenchyma; Martínez-Cabrera *et al*. 2009). Although, vessel implosion resistance has been rarely observed (Bass 1986), Hacke *et al*. (2001) argue that incipient wall break can provoke cavitation and stop vessel implosion. Regardless the mechanism linking both variables might be, the relationship of (t/b)^2^_h_ with xylem density is clearly useful since it can shed light on the ecological strategies of fossil assemblages (*e.g*., investment in support, or life history traits such as growth rates, survival, and life span, Muller-Landau 2004), besides the more obvious link to resistance to drought.

In the combined dataset, variation in VWLR was again better explained by MAP (Table 1, Figure 4). Since the dataset had 15 outliers, was not normally distributed (Shapiro-Wilk test = 0.91, P < 0.001, by group Shapiro-Wilk test 5 out of 14 communities had not normal distribution, Table S2), and the variance was not homogeneous among groups (Levene test = 3.8, P < 0.001), we performed a Welch one-way test (instead of a one-way ANOVA) to determine if there significant differences in VWLR between communities. We found significant overall differences among communities (Welch = 20.6, P < 0.001, N = 279), as well as differences between community pairs (Games-Howell test is presented in Table S3). Drier communities had vessels with thicker walls relative to their diameter (Figure 5A). These differences are in general recognized by the Games-Howell test, especially when communities on opposite sides of the MAP gradient are compared (Table S3). Despite its clear relationship with the environment, the resolution of VWLR allows to distinguish only broad MAP differences, these differences are harder to detect among communities growing at similar MAP values (*i.e.,* communities growing under a MAP below 663 mm, not significantly different among them).

**Figure 4.**
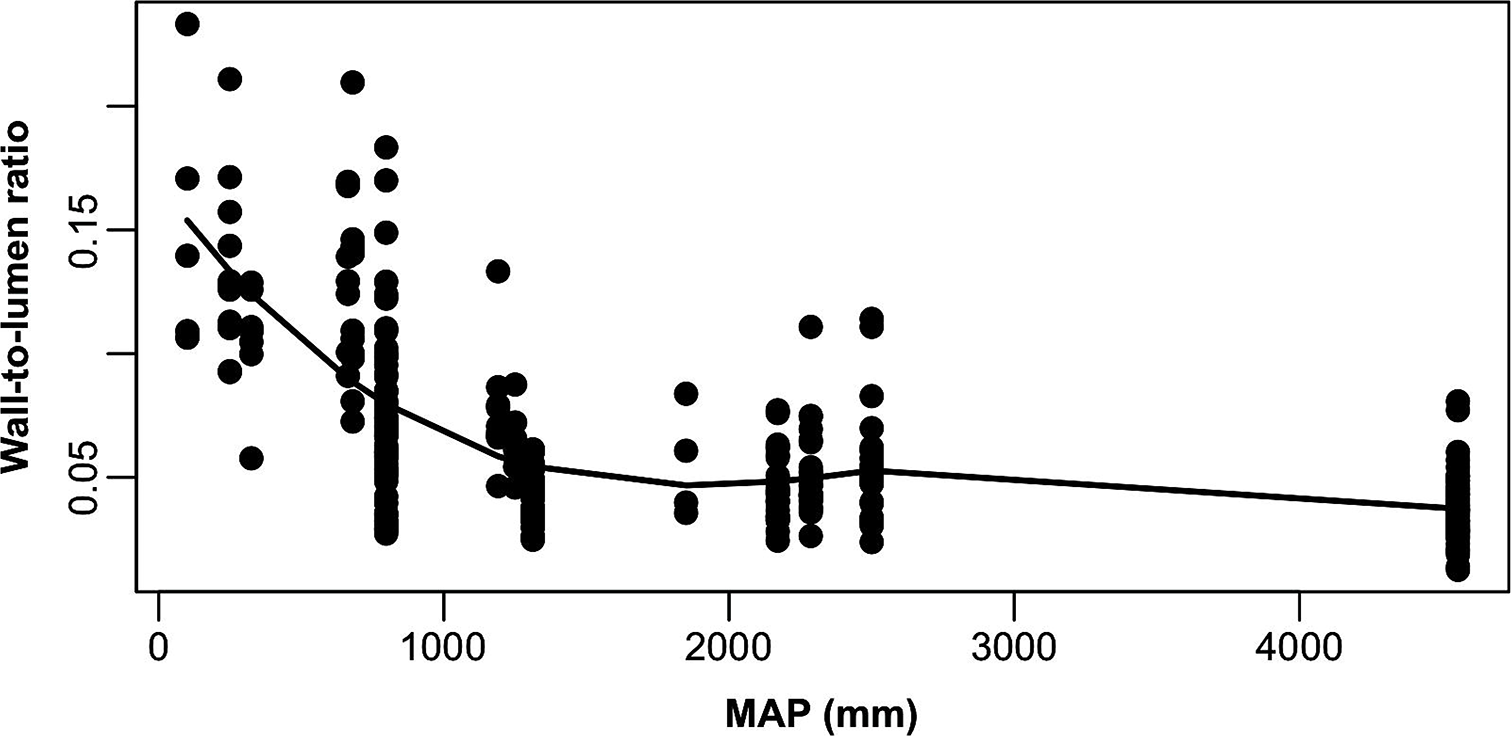
VWLR as a function of MAP.

**Figure 5.**
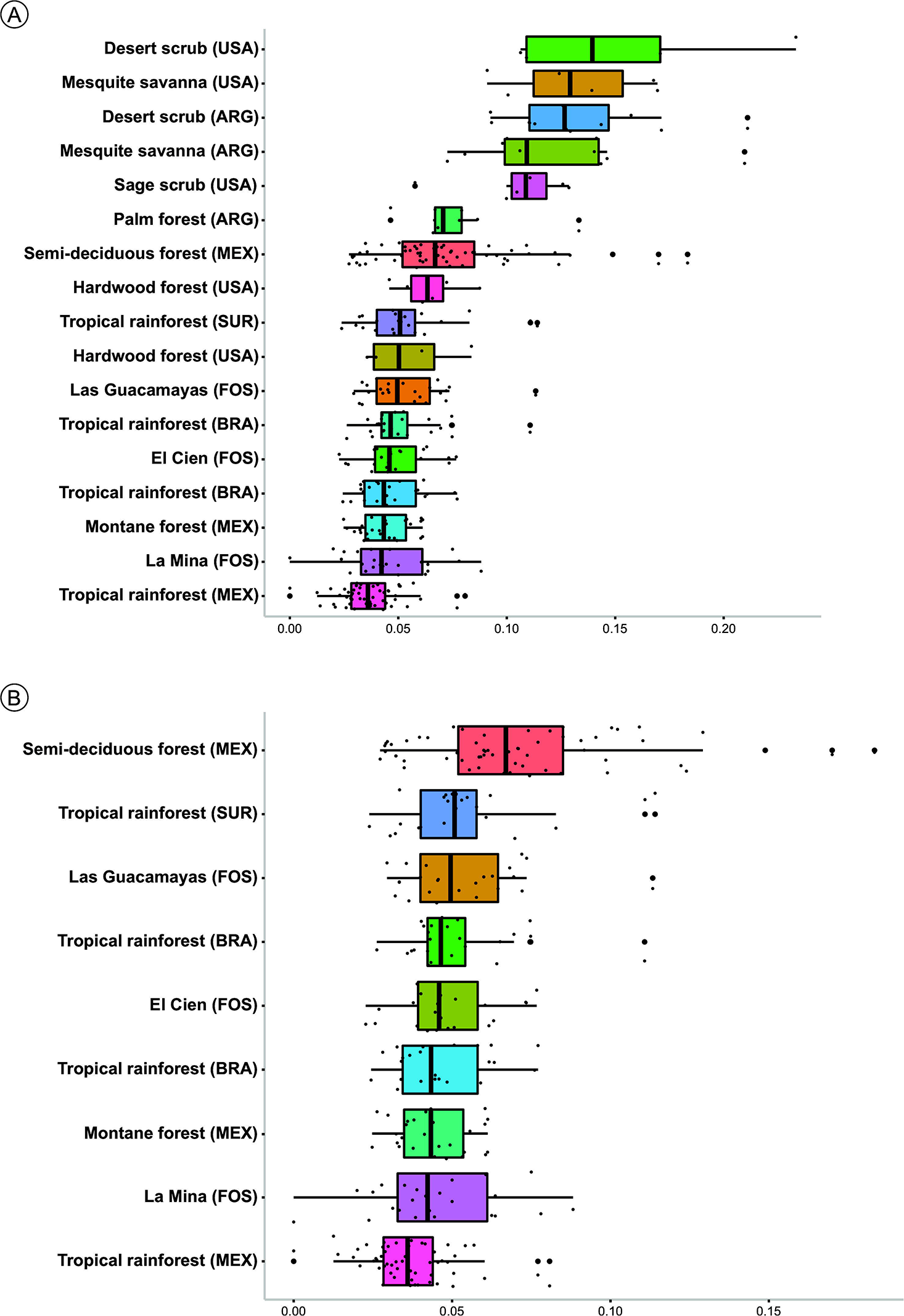
Comparison of VWLR values across extant and fossil communities. A. Complete dataset. B. Communities from seasonally dry to humid tropical forest and fossil localities. Boxplots show median, interquartile range and largest values within 1.5.times the interquartile range below the 25^th^ and above the 75^th^ percentiles. ARG=Argentina, BRA=Brazil, FOS= Fossil localities, MEX=Mexico, SUR= Suriname, USA= United States of America.

### Analysis of wall-to-lumen ratio trends in three Mexican fossil wood assemblages

All three fossil assemblages were significantly different from the drier sites (< 663 mm). From the analysis of a subset of communities only including the tropical wet/semi deciduous sites (Figure 5B; Games-Howell test Table S4) and the fossil wood localities, we found that the VWLR of Las Guacamayas (Miocene,) and El Cien Formation floras (Oligocene-Miocene) had vessels with significantly lower values of reinforcement than the semi-deciduous forest from Chamela (798 mm) and had significantly higher reinforcement values than the super humid tropical rain forest of Los Tuxtlas (4556 mm). While La Mina (Miocene, Tlaxcala) locality, was only different from the semi-deciduous forest. These results generally agree with previous paleoecological and paleoclimate analyses of those localities. Paleoclimate models suggest that La Mina and Las Guacamayas floras grew under high humidity (MAP= 2172 and 1866 mm respectively) and temperature (MAT > 25 °C), and an indistinct or short dry season, conditions typically present in tropical rain forest (Castañeda-Posadas, 2007). The case of La Mina is interesting since it has high prevalence of tropical genera such as *Terminalia*, *Cedrela* and cf. *Pterocarpus*, and together with its predicted MAP and low VWLR, not statistically different to the wettest extant tropical rain forest we analyzed, suggest low drought resistance. It is however, worth noting that the estimated wood density for this community is high (Martinez-Cabrera *et al*. 2012), which might be indicating a higher drought resistance than VWLR alone might suggest. It has been found that trees with denser wood are capable to retain more water and survive lower water potential (trees with high wood density have lower leaf turgor loss point, Fu & Meinzner 2019). This merits further research of the anatomy of this locality to evaluate if the thicker fibers (hence the high predicted wood density), might be buttressing vessels as has been hypothesized (Jacobsen *et al*. 2005), as La mina flora woods have relatively thin walls relative to their lumen (low VWLR).

El Cien Formation flora is functionally and compositionally similar to the semi-deciduous forests of the western coast of Mexico (*i.e.* Chamela). Here we found significantly lower VWLR values of El Cien Formation woods, suggesting that cavitation occurred at less negative water potential that in its living homologous Chamela. El Cien Formation flora has a strikingly similar estimated wood density to Chamela (Martínez-Cabrera *et al*. 2012), but dissimilarities in conduction efficiency and now in vessel wall reinforcement, highlight the value of incorporating more functional metrics in paleoecological analyses to recognize these nuances in fossil forest function, as was also the case for La Mina locality.

The use of the vessel wall reinforcement metric calculated with the mean hydraulic diameter ((t/b)^2^_hydraulic mean_) can be used as a sound approximation of the original (t/b)^2^_h_, if this is impractical to measure. We found that the cavitation resistance (P_50_ values) estimated with (t/b)^2^_hydraulic mean_ is close to those obtained with (t/b)^2^_h_, As the metrics have different degrees of uncertainty, we suggest avoid comparing floras using P_50_ estimates calculated with different wall reinforcement metrics. Surprisingly, VWLM was more closely related to climate, particularly to MAP, than any other anatomical trait. This suggests that these two metrics, VWLR and (t/b)^2^_h_ could be describing a slightly different ecological axis of variation, with the latter especially related to investment in support tissue in addition to water availability. Although VWLM could only be used to detect broad MAP differences among fossil assemblages, its real value lies in its use in combination with additional functional traits estimated with other anatomical variables (e.g., hydraulic conductivity, wood density) to better describe fossil forest functional strategies.

## Acknowledgments

This research was funded by Secretaría de Investigación y Posgrado – Instituto Politécnico Nacional (20220097) grant to E.E.R.

## Supplementary data

Supplemental material for this article can be accessed here: <URL added by journal>

**Table S1.** Localities and climate variables.

**Table S2.** Results of the Shapiro-Wilk test for VWLR in extant localities.

**Table S3.** Games-Howell test for the entire database results.

**Table S4.** Games-Howell test results a subset including only semi-deciduous and wet tropical communities.

